# First Lithic Age Caribbean genomes document pre-Ceramic genetic continuity and affinities to Central America and northern South America

**DOI:** 10.64898/2026.05.12.724636

**Authors:** Kendra Sirak, Adolfo José López Belando, Daniel Shelley, Manuel Garcia Arévalo, Dana Shelley, Swapan Mallick, Nadin Rohland, David Reich

## Abstract

The population history of the Caribbean’s first inhabitants has been challenging to reconstruct because few human remains are known from the region’s earliest occupation which began around 6,000 years ago in Hispaniola, Cuba, and Puerto Rico. We generated genome-wide data from 19 individuals from Hispaniola’s Samaná Peninsula and focused on four who lived during the earliest pre-Ceramic “Lithic Age”. Extending the Caribbean genetic record by more than a millennium to ∼4,400 calBP, we show that pre-Ceramic Age populations across Hispaniola and Cuba derive from a single ancestry source and document long-term genetic continuity across islands, with some local genetic structure within Hispaniola. Pre-Ceramic Age Caribbean ancestry shares most drift with populations from Central America and northern South America, although no sampled mainland group provides an adequate proxy. We infer very small effective community sizes, consistent with locally structured mating pools and little evidence of close-kin mating. These findings extend our understanding of Caribbean population history into its earliest phase.

**Teaser:** Early Caribbean people shared ancestry across islands and lived in small, locally structured communities.

## Introduction

The insular Caribbean was the final frontier settled during human migrations in the Americas, with the first evidence of occupation in Hispaniola and Cuba dating ∼6000 years Before Present (BP). A classic chronological framework for the Caribbean defines a pre-Ceramic occupation encompassing Lithic and Archaic Ages (*1*), distinguished primarily by differences in material technologies. The Lithic Age (beginning ∼6000 BP) is characterized by flaked-stone technologies, particularly blade production (*2*), and in Hispaniola and central Cuba is associated with the Casimiran (Casimiroid) series. The Archaic Age, beginning ∼4000 BP (*1, 3*) is marked by a shift toward ground-stone tool technologies and increased exploitation of marine resources, with regional variants including the Redondan Casimiroid in Cuba (∼4000-2400 BP and as late as 500 BP in some parts of the island) and the Courian Casimiroid in Hispaniola (∼4000-1800 BP (*3*)). While some traditional models associated each Age with a distinct wave of migration of people from the American continent, more recent archaeological work has questioned whether these Ages are meaningfully distinct and challenged the perception of the earliest settlers as aceramic hunter-gatherers or fisher-hunter-gatherers. A growing body of evidence documents some ceramic use (*3-8*) and incipient agriculture (*9, 10*) prior to the widespread expansion of Ceramic Age populations after ∼2500 BP who introduced traditions of intensive ceramic usage and a heavy reliance on agriculture into the Caribbean (*2, 11*). Although the archaeological record has expanded considerably, it has yet to resolve the origins of and relationships among the Caribbean’s pre-Ceramic Age inhabitants (the term “pre-Ceramic” is used here to refer collectively to people who lived before the movement of intensive ceramists/agriculturalists into the insular Caribbean ∼2500 BP).

Two key topics in research on the pre-Ceramic Age Caribbean are (1) the geographic source of the earliest migration into the Caribbean; and (2) whether cultural transitions during pre-Ceramic times reflect multiple waves of migration from the American continent. Locations as diverse as the Yucatan Peninsula (*12*), Belize (*1*), Nicaragua (*13*), northwestern South America (*2, 14-16*), or the southeast of what is now the United States (*17, 18*) have been put forward as possible locations for the earliest (Lithic Age) movement into the Caribbean. Models proposing a second movement associated with the spread of Archaic Age ground-stone technologies from northern South America place the origin of this population in northern South America (likely Venezuela via Trinidad (*19*)), while others argue for continuity across the Lithic and Archaic Ages (*20, 21*) featuring local cultural transformation without major population replacement. To date, these hypotheses have been developed primarily from material evidence due to a lack of skeletal remains from Lithic Age contexts.

Recent excavations on the Samaná Peninsula (northeastern Dominican Republic; **Supplementary Text**) provide a rare opportunity to address open questions by linking well-contextualized archaeological assemblages with the oldest human skeletal remains in the insular Caribbean to date. Regional survey identified a complex of cave and rock shelter sites within the Monumento Natural Cabo Samaná. Excavations in 2022-2023 focused on three sites within this complex – Cueva Funeraria de Daniel, Abrigo Daniel, and Abrigo Dana – as well as Playa Madama located ∼5 km north. Human skeletal remains were recovered from Cueva Funeraria de Daniel, Abrigo Daniel, and Playa Madama. Material culture across these sites, including finely finished lithic tools and evidence for diverse marine and terrestrial subsistence, is consistent with Lithic Age Casimiroid-associated occupations. The archaeological assemblages indicate expert navigation of the surrounding seas, long-distance exchange networks, the construction of wooden structures protected by rock shelters, the use of dedicated funerary spaces, and dependence on some agriculture as well as foraging for subsistence (*22*). At Cueva Funeraria de Daniel and Abrigo Daniel, dedicated funerary contexts are associated with the habitation site of Abrigo Dana (∼300 m away), suggesting structured use of the landscape. Contextual radiocarbon dates from postholes and hearths at Abrigo Dana indicate sustained pre-Ceramic Age occupation spanning at least two millennia (∼5400-3000 calBP) (*22*).

Extending the genomic record of the Caribbean by at least a millennium, we report genome-wide ancient DNA data from individuals excavated from sites on the Samaná Peninsula and integrate these data with published ancient genomes from across the Americas. These data furnish insight into the population structure and geographic affinities of the Caribbean’s earliest inhabitants.

## Results

### Ancient DNA data generation

We extracted DNA using bone or tooth samples from 36 unique individuals from three sites in Samaná (**Fig. 1B**) and generated genome-wide ancient DNA data using in-solution enrichment for ∼1.4 million single-nucleotide polymorphisms (SNPs) (**table S1**) (*23*). After filtering for samples with fewer than 20,000 SNPs or with evidence of contamination, we retained 19 individuals (**Materials and Methods**). We merged our newly generated data with published ancient and modern data from the Americas, restricting to the SNPs on the “Compatibility Panel” which has been optimized to minimize technology-specific biases (*24*). Central to our analyses are published reference data from the pre-Ceramic Age Caribbean, including an individual from southeastern Hispaniola (*DominicanRepublic_Andres_PreCeramic*, hereafter “*DA*”), dated to 3141-2954 calBP (2890 ± 20 BP, PSUAMS-7291), which places this individual as living in the Caribbean’s Archaic Age, as well as a genetically homogeneous group of 50 Archaic Age individuals from Cuba (*Cuba_PreCeramic*, hereafter “*CA*”) who lived between ∼3300-600 calBP (**Fig. 1A**).

**Fig. 1.**
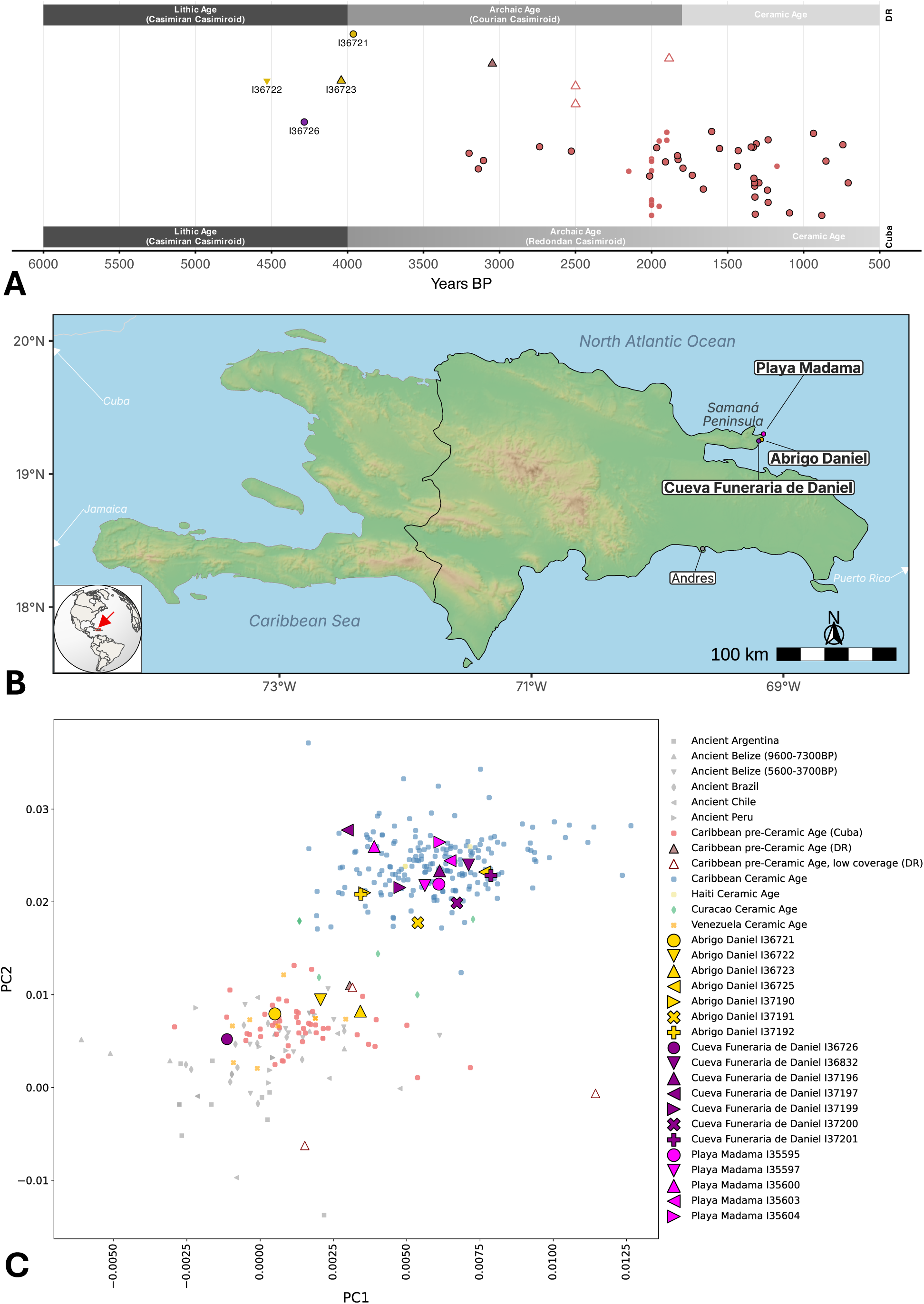
Temporal and genetic overview of pre-Ceramic and Ceramic Age Caribbean data. (**A**) Chronological placement of pre-Ceramic Age individuals from the Dominican Republic (DR) and Cuba. Points are plotted by median calendar year BP. The dates of points outlined in black were determined through direct ^14^C dates on bone, while non-outlined points indicate context dates. Point color and shape identical to Panel C. The upper and lower bars show cultural-historical periodizations used for the DR and Cuba, respectively. Four labeled individuals from Samaná (I36721, I36722, I36723, and I36726) are highlighted. (**B**) Map of Hispaniola showing the locations of archaeological sites on the Samaná Peninsula analyzed in this study. Colored points mark sites with samples newly reported in this work, including Playa Madama, Abrigo Daniel, and Cueva Funeraria de Daniel; Andrés is shown for reference. The inset map shows the location of Hispaniola within the Caribbean. (**C**) PCA shows a pre-Ceramic Age-associated genetic signature for four Samaná individuals and a Ceramic Age-associated genetic signature for 15.

### Chronology and archaeology

We generated direct ^14^C dates on bone for 15 unique individuals from Samaná for whom we also report DNA (**table S1** and **S2**; **Supplementary Text**). Twelve dates fall within the Ceramic Age, while I37621 and I36723 from Abrigo Daniel and I36726 from Cueva Funeraria are dated to the Lithic Age (*1*) in line with expectations from archaeological material (*25*). I36726, the oldest individual in our dataset, is directly dated to 3870 ± 30 BP (PSUAMS-14458; calibrated to 4411– 4158 calBP without a marine reservoir correction or 4342–3930 calBP under a mixed marine– terrestrial model with a Samaná peninsula-specific correction (*26*); **table S2**). This makes I36726 more than 1,000 years older than the previously oldest reported Caribbean genome (CAO019 from Canímar Abajo in Cuba was dated to 3324-3077 calBP (3000 ± 19 BP, MAMS-44074)) (*27*). I37621 and I37623 from Abrigo Daniel date to 4079–3845 calBP (3630 ± 30 BP, Beta-765015) and 4150–3933 calBP (371 ± 30 BP, Beta-765018), respectively, placing them several centuries later than I36726, and presenting an opportunity to analyze time-transect data within Samaná, across Hispaniola, and between Hispaniola and Cuba.

I36726 from Cueva Funeraria was interred in a supine position with evidence suggesting that the body was wrapped at the time of burial; this burial style was distinct from that observed in individuals from the same site directly dated to the Ceramic Age. Three individuals from Abrigo Daniel (I36721, I36722, and I36723) were also buried in a supine position (with I36721 and I36722 also wrapped) to the exclusion of other individuals from that site who dated to the Ceramic Age. While I36721 and I36723 were radiocarbon dated, we did not obtain a date from I36722; however, this individual was associated with a date on charcoal of 4530 ± 30 BP (Beta-638468). The correlation between burial style and Lithic Age radiocarbon dates, as well as concordance with genetic clustering patterns described below, provides further support for a Lithic Age date for I36722.

### Genetic homogeneity in the pre-Ceramic Age Caribbean

We first used Principal Component Analysis (PCA) to visualize the genetic structure of the ancient individuals from Samaná relative to published individuals from the Americas, computing the axes from a set of reference groups that maximally differentiates ancestry within the Caribbean (**Fig. 1C**). In this PCA, individuals from the Ceramic Age Caribbean form a genetic cluster, whereas pre-Ceramic Age individuals cluster separately near to other ancient Americans. Projecting our new data onto this PCA, four individuals from Samaná – one from Cueva Funeraria de Daniel and three from Abrigo Daniel – fall into the Caribbean pre-Ceramic Age cluster. This is supported by *f*_*4*_ statistics showing excess allele sharing with previously published Caribbean pre-Ceramic Age individuals (|Z| > 3) (**table S3**). The sites of Cueva Funeraria and Abrigo Daniel both contain pre-Ceramic occupation layers characterized by Lithic Age “Samaneses de tradición Casimiroide” artifact styles (*1*). The remaining 15 individuals – six from Cueva Funeraria, four from Abrigo Daniel, and all five from Playa Madama – fall within the Caribbean Ceramic Age cluster. Notably, while all Playa Madama individuals have Ceramic Age-associated ancestry with the oldest directly dated to 1415 ± 20 BP (PSUAMS-14876), an older date on charcoal from the site (2795 ± 140 BP, I-9780) and the absence of ceramic assemblages (*28*) suggests that the site was in use long before the lifetimes of the individuals analyzed here. The four Samaná individuals with pre-Ceramic Age genome-wide genetic signatures belong to mitochondrial DNA (mtDNA) haplogroup D1 (**table S1**), the most common haplogroup in the pre-Ceramic Age Caribbean. Haplogroup D1 decreased in frequency during the Ceramic Age, concurrent with an increase in C1 and A2 lineages common in the Caribbean today (*27, 29-32*).

In what follows, we focus specifically on resolving the genetic relationships of the four individuals from the Samaná Peninsula with pre-Ceramic Age signatures.

### Population structure in pre-Ceramic Age Samaná

To investigate genetic relationships within pre-Ceramic Age Samaná, we tested for population structure among the sampled individuals. *f*_*4*_-statistics reveal internal structure among the four individuals with pre-Ceramic Age genetic signatures, indicating that I36722 and I36723, both sub-adult males buried near each other at Abrigo Daniel form a clade relative to I36721 (a male from Abrigo Daniel) and I36726 (a male from Cueva Funeraria) (|Z| > 8.2) (**Table 1**; **table S4**). As I36722 and I36723 show slightly elevated contamination estimates and lower data quality relative to I36721 and I36726 (**table S1**), we repeated tests using damage-restricted versions of these samples (**Materials and Methods**). We find that the strong affinity between I36722 and I36723 is robust to damage-restriction and persists across multiple data subsets, indicating that it is not driven by post-mortem damage or library-specific artifacts. This supports the presence of a genuine affinity between these individuals, and we merge I36722 and I36723 as *Samaná_PreCeramic_AD* (‘*AD*’ for Abrigo Daniel) in some subsequent analyses. There is no evidence for structure among the remaining individuals (**table S4**).

**Table 1.** Select statistics of the form *f*_*4*_(*Outgroup, Samana1*; *Samana2, Samana3*) show significantly greater allele sharing between I36722 and I36723 relative to I36721 and I36726. Columns report Z-scores for three formulations of the *f*_*4*_-statistic test. Results are shown for both full and damage-restricted (‘_d’ suffix) data. SNPs indicate the number of sites used for each test. Significant results (|Z| > 3) in bold. All data are in **table S4**.

| Outgroup | A | B | C | $f_4(\text{Out}, \text{A}; \text{B}, \text{C})$ | $f_4(\text{Out}, \text{B}; \text{A}, \text{C})$ | $f_4(\text{Out}, \text{C}; \text{A}, \text{B})$ | SNPs |
| --- | --- | --- | --- | --- | --- | --- | --- |
| Cameroon_SMA | I36723 | I36722 | I36721 | **-9.993** | **-8.188** | 2.096 | 681904 |
| Cameroon_SMA | I36723 | I36722_d | I36721 | **-9.355** | **-8.346** | 1.075 | 412551 |
| Cameroon_SMA | I36723_d | I36722 | I36721 | **-8.243** | **-6.756** | 1.597 | 261195 |
| Cameroon_SMA | I36723_d | I36722_d | I36721 | **-7.642** | **-6.655** | 0.896 | 167151 |
| Cameroon_SMA | I36723 | I36722 | I36726 | **-10.765** | **-10.615** | -0.226 | 686609 |
| Cameroon_SMA | I36723 | I36722_d | I36726 | **-9.922** | **-9.947** | 0.259 | 414063 |
| Cameroon_SMA | I36723_d | I36722 | I36726 | **-9.326** | **-8.786** | -0.616 | 262147 |
| Cameroon_SMA | I36723_d | I36722_d | I36726 | **-8.709** | **-8.47** | -0.195 | 167452 |
| Cameroon_SMA | I36726 | I36721 | I36723 | 0.805 | 1.345 | 0.471 | 699561 |
| Cameroon_SMA | I36726 | I36721 | I36722 | 0.979 | 2.70 | 1.702 | 1066122 |

### Shared ancestry across the pre-Ceramic Age Caribbean

To test for population structure in the pre-Ceramic Age Caribbean, we computed *f*_*4*_(*Outgroup, Caribbean_PreCeramic1; Caribbean_PreCeramic2, Caribbean_PreCeramic3*), rotating *Cuba_PreCeramic* (‘*CA*’), *DominicanRepublic_Andres_PreCeramic* (‘*DA*’), *Samaná_PreCeramic_AD*, I36721, and I36726 through each position (**table S5**). All individuals from Samaná are consistent with deriving from a single ancestry source relative to CA and DA, as statistics of the form *f*_*4*_(*Outgroup, CA/DA; Samana1, Samana2*) do not deviate significantly from zero. Comparisons among pre-Ceramic Age individuals from Cuba, Andrés, and Samaná are also non-significant, indicating symmetrical relatedness. Together, these results document shared ancestry among pre-Ceramic Age people across Hispaniola and Cuba spanning nearly four millennia from at least 4411–4158 calBP (I36726, 3870 ± 30 BP, PSUAMS-14458) to 735–679 calBP (CAO028, 812 ± 20 BP, MAMS-44083 (*27*)). This is consistent with a single population movement into the Caribbean followed by limited divergence, such that genetic drift has not produced a clear phylogenetic separation among island populations.

Despite shared ancestry and signal of inter-island symmetry, fine-scale analyses uncover some internal structure within Hispaniola. All people from Samaná share significantly more alleles with each other relative to *CA* (|Z|=2.7-9.1); however, only I36721 and *Samana_PreCeramic_AD* share significantly more alleles relative to *DA* (|Z|=3.4-5.1) while I36726 (the oldest directly dated individual in our dataset) is symmetrically related to I36721, *Samana_PreCeramic_AD*, and DA. This pattern defines internal structure within Samaná, with I36721 and *Samana_PreCeramic_AD* – all from the site of Abrigo Daniel – forming a closer genetic unit relative to I36726 from Cueva Funeraria.

Pairwise *qpWave* analyses are consistent with all pre-Ceramic Age individuals from Samaná, Cuba, and Andrés deriving from a single ancestry source relative to a diverse reference set of published ancient and modern Americans (p > 0.05; **table S6**), a pattern that also holds when all are tested jointly (p=0.23). Using a rotating *qpWave* strategy, models are not rejected when *Cuba_PreCeramic* or *DominicanRepublic_Andres_PreCeramic* is included in the reference set, indicating no differential relatedness at the inter-island scale. However, when *Samaná_PreCeramic_AD* is added to the reference set, symmetry between Samaná individuals and either Cuba or Andrés is strongly rejected (p < 10^−4^), while symmetry within Samaná and between Cuba and Andrés is maintained. Together, these results indicate that pre-Ceramic Caribbean populations are genetically homogeneous at a broad scale, arguing against population replacement from a mainland source at the transition from the Lithic to Archaic Age in Hispaniola. Nevertheless, individuals from Samaná share excess genetic drift relative to Cuba and Andrés, revealing subtle population structure within an otherwise largely uniform pre-Ceramic Age Caribbean gene pool.

To further test the single-ancestry model for all pre-Ceramic Age Caribbean people inferred through *f*_*4*_-statistics and *qpWave* and attempt to refine the geographic source or sources of the early settlers of the Caribbean (although inference is constrained by the relative sparsity of published ancient DNA data from the Americas), we looked for asymmetric relationships between diverse ancient populations from the American continent plus the Ceramic Age Caribbean (*Ancient_American*) and any pair of *Cuba_PreCeramic, DominicanRepublic_Andres_PreCeramic, Samana_PreCeramic_AD*, I36721, and I36726 using the statistic *f*_*4*_*(Outgroup, Ancient_American; Caribbean_PreCeramic1, Caribbean_PreCeramic2)* (**table S7**). *f*_*4*_-statistics were overwhelmingly consistent with symmetry, with no *Ancient_American* population showing a consistent excess affinity to any pre-Ceramic Age Caribbean group, supporting a model of shared ancestry across the pre-Ceramic Age Caribbean rather than distinct mainland-derived ancestry components. A parsimonious explanation of the data is that a migration to Hispaniola and Cuba during the Lithic Age established a pre-Ceramic Age population that experienced no substantial gene flow from genetically distinct groups from the American mainland until the major biological and cultural shifts associated with the onset of the Ceramic Age (for caveats, see Discussion). Based on evidence of relative homogeneity and to increase statistical power, we pooled all pre-Ceramic Age Caribbean individuals from Hispaniola and Cuba as a single group (*Caribbean_PreCeramic*) and tested its relationship to continental populations using *f*_*4*_*-*statistics.

### Pre-Ceramic Age Caribbean ancestry shares most drift with populations from the Isthmo-Colombian region and northern South America

The ancestry of *Caribbean_PreCeramic* is consistent with deriving wholly from the expansion of Southern Native American (SNA) lineages into South America with no evidence of admixture from other genetic lineages, such as Anzick-1-related ancestry that spread into Central and South America via subsequent migration events (**table S8**) (*33, 34*). We therefore focused on resolving the relationship between *Caribbean_PreCeramic* and other groups with SNA-related ancestry. To broadly visualize shared genetic drift among these populations, we first computed pairwise outgroup-*f*3 statistics of the form *f*3(*Outgroup; Ancient_American1, Ancient_American2*) and used these values to construct a neighbor-joining tree (**Fig. 2B**; **table S9**). In this tree, *Caribbean_PreCeramic* falls within a broader grouping that includes *Caribbean_Ceramic* as well as ancient populations from the Isthmo-Colombian region (Panama), northern South America (Venezuela), and Mexico’s Yucatán Peninsula, consistent with elevated shared drift with these regions. By contrast, populations from Brazil, the Andes, and the Southern Cone occupy more distant branches of the tree, consistent with lower levels of shared drift with *Caribbean_PreCeramic*.

**Fig. 2.**
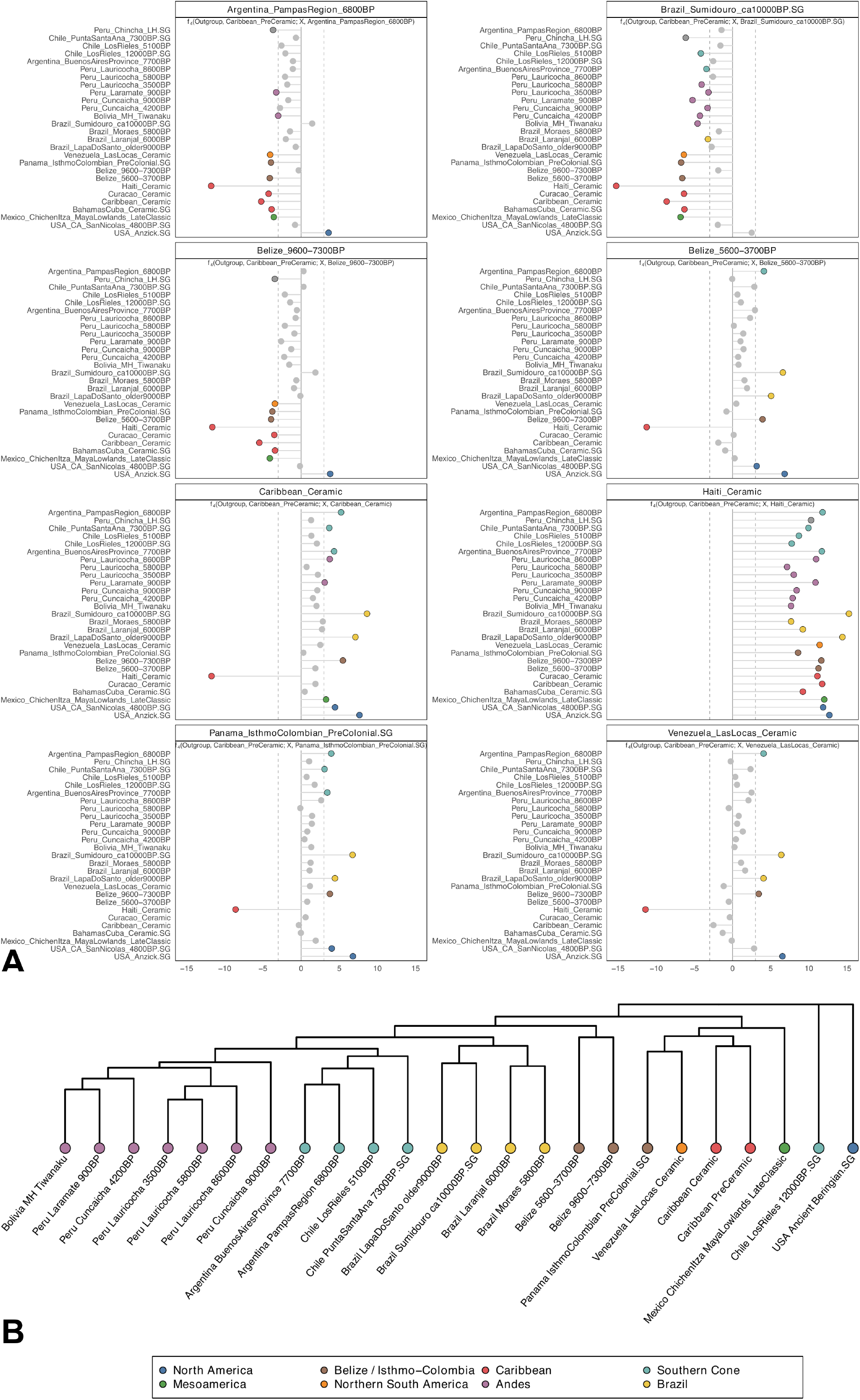
Shared drift and allele-sharing patterns involving *Caribbean_PreCeramic*. (**A**) *f*_4_-statistics showing asymmetric patterns of allele sharing between *Caribbean_PreCeramic* and eight focal ancient American populations. Each panel shows statistics of the form *f*_*4*_(*Outgroup, Caribbean_PreCeramic*; *X, Focal_Ancient_American*). Positive Z-scores indicate greater allele sharing between *Caribbean_PreCeramic* and the focal population (listed at the top of each panel), and negative Z-scores indicate greater allele sharing with the comparison population (listed along the left side of each panel). Points represent Z-scores with dashed vertical lines marking the significance threshold (|Z|=3). Points with |Z| < 3 are shown in gray, while significant values are colored by the geographic affiliation of the comparison population (North America, Mesoamerica, Belize/Isthmo-Colombian region, northern South America, Andes, Southern Cone, Brazil, and Caribbean; legend on bottom). Data are in **table S8**. (**B**) Neighbor-joining tree based on pairwise outgroup-*f*_3_ values among select ancient American populations; data are in **table S9**. Outgroup-*f*_3_ values were converted to distances as 1/*f*_3_ for tree construction. The tree is rooted for visualization with *USA_Ancient_Beringian*.*SG*. Because pairwise outgroup *f*_3_ values summarize shared post-outgroup drift, this tree is interpreted as a visualization of shared-drift affinities rather than a uniquely resolved phylogeny. Branch lengths were rescaled for visualization, and terminal branches were equalized for display. Populations are colored by geographic affiliation. Legend on bottom applies to both panels.

We use statistics of the form *f*_*4*_(*Outgroup, Caribbean_PreCeramic; Ancient_American1, Ancient_American2*) to provide finer-scale resolution of patterns of shared drift, documenting structure in the relationships of *Caribbean_PreCeramic* to ancient continental American and later Ceramic Age Caribbean groups (**table S10**). We find that *Caribbean_PreCeramic* shares excess alleles with ancient populations from the Ceramic Age Caribbean, northern South America, and the Isthmo-Colombian region relative to more southerly (for example, Argentina Pampas 6800BP (*33*)) or highly drifted early diverging lineages (for example, Brazil Sumidouro (*35*)) (**Fig. 2A**), consistent with its placement in the neighbor-joining tree. Excess allele sharing between pre-Ceramic Age and Ceramic Age Caribbean groups likely reflects both deep shared ancestry and direct admixture from pre-Ceramic Age groups into the Ceramic Age gene pool (ranging from ∼2% in *Caribbean_Ceramic* to ∼18% in *Haiti_Ceramic*). Consistent with this, *Haiti_Ceramic* (which harbors the largest ancestry contribution from *Caribbean_PreCeramic*) shares significantly more alleles with *Caribbean_PreCeramic* than do other Ceramic Age people from the insular Caribbean (|Z| = 11.7). Beyond the Caribbean, the observed affinity to populations from the Isthmo-Colombian region and northern South America indicates a relationship to lineages from these regions but affirm that no sampled mainland population provides an adequate proxy for its ancestry.

While *f*_*4*_-statistics do not resolve a geographic source for pre-Ceramic Age Caribbean ancestry, some targeted comparisons provide additional insight into its relationship to mainland populations. In particular, comparisons involving two temporally distinct ancient populations from Belize highlight informative differences. Previous work has modeled the later *Belize_5600–3700BP* population as a mixture of (1) an earlier group, *Belize_9600–7300BP*, representing a deep lineage associated with early southward dispersals into the Americas, and (2) a lineage related to present-day Chibchan-speaking populations in southern Central America and northwestern South America (*36*). We find that *Caribbean_PreCeramic* and *Belize_9600–7300BP* both share significantly more alleles with *Belize_5600–3700BP*, while *Belize_5600–3700BP* is not significantly closer to either (**Table 2**). This pattern is consistent with the broader signal observed in *f*_*4*_-statistics, in which *Caribbean_PreCeramic* shows relatively greater affinity to populations from the Isthmo-Colombian region and northern South America than to other lineages. Importantly, this pattern of allele-sharing with the earlier and later Belize groups is not unique to *Caribbean_PreCeramic*, but is also observed in Ceramic Age Caribbean groups (*29*), Late Classic Maya from the Yucatán Peninsula (*37*), pre-Columbian Panamanians (*38*), and ceramic users from Venezuela older than ∼2100 calBP (*29*) (**table S11**). This supports a broad geographic distribution of this ancestry signal across Central America and northern South America.

**Table 2.**
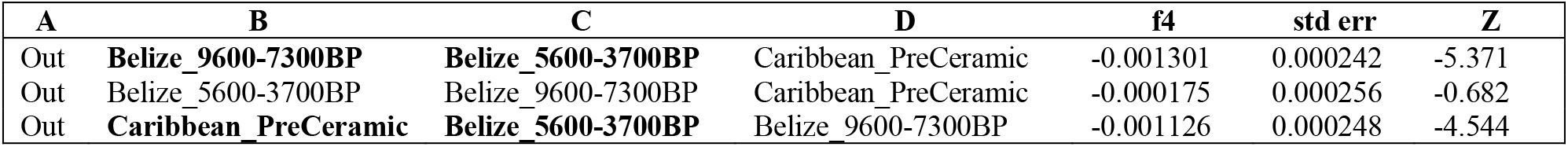
*f*_*4*_-statistics testing allele-sharing relationships among *Caribbean_PreCeramic* and two ancient Belize populations. Significantly negative values (|Z| > 3) indicate excess allele sharing between *Belize_5600–3700BP* and both *Caribbean_PreCeramic* and *Belize_9600– 7300BP*, while the non-significant result in the second row shows that *Belize_5600–3700BP* is symmetrically related to the other two. Standard errors and Z-scores are shown for each test.

To further investigate the phylogenetic relationships among populations exhibiting this pattern, we used *qpWave* to test whether they could be modeled as deriving from a single ancestry stream relative to a set of reference populations. Pairwise tests show that *Caribbean_PreCeramic* forms a clade with ancient Panamanians and Venezuelan populations (p = 0.28 and 0.69, respectively), but clade models are rejected for *Caribbean_PreCeramic* with Maya from the Yucatán or Ceramic Age Caribbean groups (p < 0.01; **table S12**). However, when any of these populations is included in the reference set, models of a shared ancestry stream are rejected, indicating that their relationships are not consistent with a simple tree-like structure. Instead, these results support a model of more complex population structure, potentially involving descent from a common but unsampled ancestral population followed by regional differentiation following dispersal.

### *qpGraph* supports multiple models of *Caribbean_PreCeramic* ancestry

We finally fitted admixture graphs using *qpGraph*, increasing model complexity while always allowing zero to two admixture events, and assessing model fit using the maximum absolute Z-score of residuals. We first fit a simplified admixture graph including only six populations chosen to capture major deep structure in the Americas (**fig. S1**). In this reduced set, an admixture-free model provides an adequate fit for the data, placing *Caribbean_PreCeramic* as a distinct lineage within the broader SNA-related variation (recapitulating the known “star-like” phylogeny of SNA ancestry (*33*)). However, this model reflects a coarse approximation, as it does not include populations that may share finer-scale ancestry with *Caribbean_PreCeramic*.

When additional populations from Central America and northern South America are incorporated, models without admixture and with a single admixture event are no longer sufficient (|Z| > 3), indicating that the simple tree structure is not consistently supported when resolution is increased. This suggests that the basal placement of *Caribbean_PreCeramic* in reduced models is not a uniquely supported history, but rather a consequence of limited sampling. Allowing two admixture events yields multiple well-fitting models (|Z| < 2.5), which can be grouped into several recurring classes of topologies (**Fig. 3**). Notably, these models differ in how they accommodate the signal of shared ancestry between *Caribbean_PreCeramic* and mainland populations. In one class of models, *Caribbean_PreCeramic* retains a basal position as an unadmixed lineage (**Fig. 3A**), with the admixture occurring among mainland populations. In contrast, other models instead place admixture directly in *Caribbean_PreCeramic* (**Fig. 3B**), modeling it as deriving ancestry from two distinct mainland-related sources. Additional models introduce deeper or alternative structures that also provide acceptable fits (**Fig. 3C**).

**Fig. 3.**
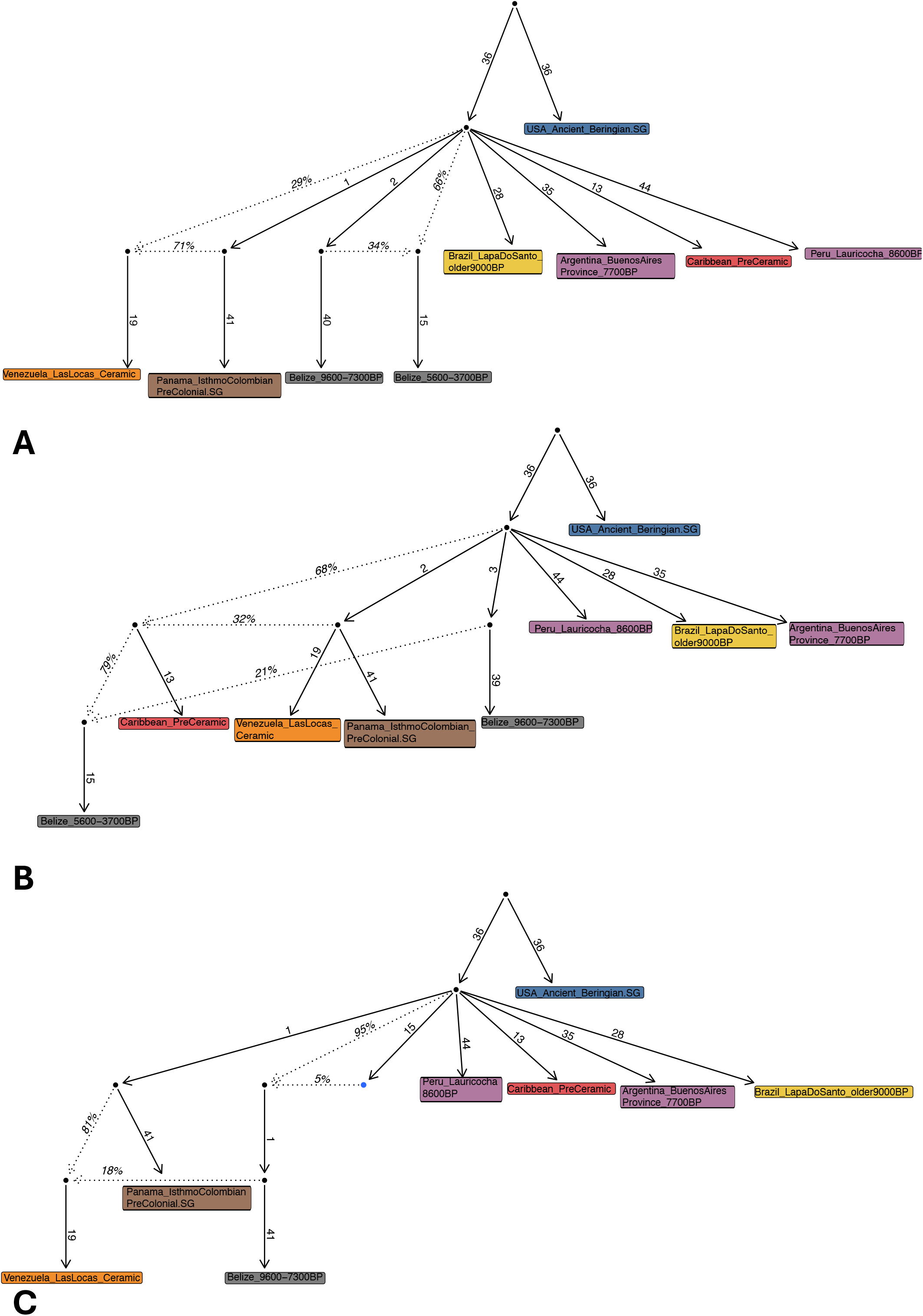
Representative *qpGraph* models illustrating alternative placements of *Caribbean_PreCeramic* ancestry. We show three well-fitting admixture graph models (|Z| < 3) that capture distinct classes of solutions. Very short branches (drift < 0.001) are collapsed for visualization. (**A**) A model in which *Caribbean_PreCeramic* is represented as an unadmixed lineage, occupying a relatively basal position within SNA-related variation, with admixture occurring among mainland populations (score = 13.6; |Z|_max_ = 2.1). (**B**) A model in which *Caribbean_PreCeramic* is admixed, indicating that the observed allele-sharing patterns can alternatively be explained by placing ancestry mixture directly in the lineage leading to pre-Ceramic Age Caribbean groups (score = 15.0; |Z|_max_ = 2.1). (**C**) A model invoking a deeper or alternative ancestry configuration that also provides an acceptable fit (score = 15.9; |Z|_max_ = 2.3). *Belize_5600– 3700BP* is connected by a near-zero drift edge and is therefore not shown as a separate terminal node, but its inferred position is indicated by a blue node.

Despite these differences in topology, all well-fitting models share a key feature: *Caribbean_PreCeramic* does not form a simple clade with any single sampled mainland population. Instead, the signal of shared ancestry with populations from Central America and northern South America can be accommodated in multiple, equally plausible ways. The existence of these alternative solutions highlights the non-uniqueness of *qpGraph* models and indicates that the ancestry of pre-Ceramic Age Caribbean populations is not adequately represented by any population that has been sampled at present. Rather, these results are best explained by descent from a structured or unsampled ancestral lineage, followed by isolation and genetic drift after dispersal into the Caribbean.

### Very small effective community sizes in Samaná

We inferred runs of homozygosity (ROH) for the four individuals from Samaná to characterize patterns of parental relatedness and background demography (*39*). Samaná individuals exhibit total ROH across all length classes at the high end of the range observed among previously studied pre-Ceramic Age Caribbean individuals (**fig. S2**) (*29*). In particular, high levels of short ROH (4-8 cM) indicate elevated background relatedness, consistent with long-term residence in a limited mating pool, while substantial intermediate ROH (8-20 cM) further support residence within a relatively small and structured population (**table S13**). No individual from Samaná exhibits sufficient long ROH (>20 cM) to be consistent with offspring of extremely close kin (e.g., first cousins). However, three individuals show moderate amounts of long ROH, consistent with parental relatedness at the level of second- or third-degree relatives. This pattern is most parsimoniously explained by demographic constraints in small communities, where the probability of mating among more distant relatives is elevated, rather than systematic close-kin unions. In contrast, I36723 shows no ROH >20 cM and a maximum segment length of 16.5 cM, indicating no detectable recent close-kin union. Together, these results suggest that while unions among more distant relatives occurred, they were not ubiquitous.

To quantify background demography, we inferred effective community size (*Ne*) from the distribution of ROH lengths (4-20 cM), which reflect co-ancestry within approximately the past 50 generations. We estimate the recent *Ne* to be 114-163 (95% confidence interval fitted via the likelihood profile) indicating a very small local breeding population. This value is lower than previously reported estimates for pre-Ceramic Age Caribbean populations (Ne=200-300) (*29*). Taken together, the combination of elevated short and intermediate ROH with only moderate levels of long ROH indicates that Samaná individuals derived from a persistently small and locally structured mating pool, in which high background relatedness accumulated over time despite the absence of systematic close-kin mating.

## Discussion

Recent studies of genome-wide ancient DNA (*27, 29*) have provided an important foundation for understanding the origins and affinities of pre-Ceramic Age Caribbean populations, although the earliest settlers have until now been characterized largely through material culture rather than biological evidence. By generating genome-wide ancient DNA data from some of the earliest individuals in the Caribbean and co-analyzing them with people who lived a millennium or more later on the islands of Hispaniola and Cuba, we provide new insight into the ancestry, population structure, and demographic history of the region’s first inhabitants.

First, we identify subtle but consistent population structure within Hispaniola. Individuals from Samaná share excess drift relative to pre-Ceramic Age individuals from southeastern Hispaniola (Andrés) and Cuba, indicating local differentiation despite overall genetic homogeneity across the Caribbean. Within Samaná, we also detect fine-scale structure, with individuals from Abrigo Daniel forming a closer genetic unit relative to an individual from Cueva Funeraria. These findings suggest structure at local scales likely reflecting limited gene flow among communities.

Second, we show that pre-Ceramic Age groups from Hispaniola and Cuba, spanning nearly four millennia, are consistent with deriving from a single ancestry source. Despite variability in material culture across the Lithic and Archaic Ages, we find no evidence for major population replacement or large-scale gene flow from distinct mainland populations during this period. This supports models of continuity across the pre-Ceramic Age sequence, in which cultural changes reflect local developments or limited interactions rather than migrations (*40*). A limitation of our sampling is that it does not include individuals associated with the “Ortoiroid” tradition, which has been hypothesized to represent a second migration from northern South America. Additional ancient DNA from Ortoiroid-associated contexts, which are found primarily but not exclusively in Puerto Rico, as well as broader sampling from the mainland, will be necessary to evaluate this hypothesis.

Third, we show that pre-Ceramic Age Caribbean ancestry derives from the broader Southern Native American (SNA) lineage and shares most drift with populations from the Isthmo-Colombian region and northern South America. This indicates that the ancestry contributing to early Caribbean populations was part of a widely distributed gene pool spanning Central America and northern South America prior to its dispersal into the Caribbean. Consistent with this, *f*_4_-statistics reveal excess allele sharing with populations from these regions, with affinities broadly distributed rather than confined to any single group for which we have genomic data. Together, these results suggest that Caribbean pre-Ceramic Age populations descended from a lineage related to, but distinct from, those in the Isthmo-Colombian and northern South American regions, with divergence likely predating the full development of regional genetic substructure on the mainland. This interpretation is consistent with archaeological and linguistic evidence pointing to southern Central America or northwestern South America as a plausible region for early Caribbean dispersals, including the diversification and spread of Chibchan languages (*41, 42*) estimated to have occurred around the time of the early movements into the Caribbean and the movement of cultivars from the Isthmo-Colombian area into the insular Caribbean (*2, 43*). However, as our genetic results do not support a simple phylogenetic placement of *Caribbean_PreCeramic* within any single known mainland lineage, we caution against equating this ancestry directly with proto-Chibchan populations.

Fourth, our *qpGraph* analyses highlight that the ancestry of pre-Ceramic Age Caribbean populations cannot be captured by a single, uniquely resolved model. While a simplified graph including only deeply divergent populations places *Caribbean_PreCeramic* as a basal lineage within SNA-related variation, this placement breaks down when additional populations from Central America and northern South America are incorporated. In more complete models, multiple well-fitting solutions emerge. In some, *Caribbean_PreCeramic* remains a basal lineage, while in others, *Caribbean_PreCeramic* is modeled as admixed. The existence of distinct but equally plausible topologies indicates that the ancestry of *Caribbean_PreCeramic* is best explained as deriving from a structured or unsampled ancestral lineage. Importantly, the appearance of *Caribbean_PreCeramic* as a basal lineage in simplified models should therefore be interpreted as an approximation under limited sampling rather than a uniquely supported historical scenario.

Ancient DNA offers a powerful window into Caribbean population dynamics, and each additional genome increases our understanding of human history in this region. Our results show that the earliest Caribbean populations were shaped by broader population processes across southern Central America and northwestern South America, advancing a framework for interpreting the Caribbean as part of a wider interconnected American landscape.

## Materials and Methods

### Permissions

We acknowledge the ancient individuals whose remains we study, present-day people who have an Indigenous legacy, and Caribbean-based colleagues centrally involved in this work. The Guahayona Institute, the group which led the excavations of the ancient individuals reported in this manuscript, works together with entities both within and outside of the Dominican Republic. This includes the García Arévalo Foundation, the Museo del Hombre Dominicano, the General Direction of Museums, the Vice-Ministry of Protected Areas, the University of Puerto Rico, the Natural History Museum of Los Angeles County, and the University of Winnipeg. Permits for archaeological excavation of the sites on the Samaná Peninsula included in this work were issued to Adolfo José López Belando by the Museo del Hombre Dominicano following an application which was approved by the Ministerio de Medio Ambiente y Recursos Naturales. This permission explicitly specified ancient DNA analysis and the generation of radiocarbon dates. Remains were handled with respect, and damage to them was minimized as much as possible, for example, by using disarticulated petrous fragments or minimally-invasive sampling techniques (*44*).

### Ancient DNA data generation

In clean-room facilities at Harvard Medical School, we generated powder from 38 tooth or cochlea samples which we assessed as representing 36 unique individuals. A total of 13 failed nuclear capture (**table S1**), leaving 23 individuals with genome-wide data. We extracted DNA following a published protocol (*45*). From the DNA extracts, we prepared truncated dual-barcoded double-stranded libraries (*46*) or dual-barcoded single-stranded libraries (*47*). For double-stranded libraries, we used a partial uracil-DNA-glycosylase (UDG) preparation (*46*), which leaves a reduced damage signal at both terminal ends of the DNA molecule (5’C-to-T, 3’G-to-A). For single-stranded libraries, we used *Escherichia coli* UDG (“USER” from NEB), which inefficiently cuts terminal and penultimate uracils. We added two seven-base-pair (bp) indexing barcodes to the truncated adapters of each double-stranded library and completed the adapter sites for sequencing (it is not necessary to add these barcodes to single-stranded libraries, which are already indexed and adapter sites are completed from library preparation) and sequenced libraries using an Illumina NovaSeq S4 flowcell or a HiSeqX10 instrument with 2 × 101 cycles to generate paired-end reads. Indices were read with 2 × 7 cycles (double-stranded libraries) or 2 × 8 cycles (single-stranded libraries). We prepared multiple libraries for some individuals to increase coverage (**table S14**).

To generate SNP capture data, we performed in-solution target hybridization using the “Twist Ancient DNA” assay (Twist Bioscience) with mitochondrial probes spiked in. The Twist reagent targets around 1.4 million SNPs, including all in the “1240k” capture set commonly used in ancient DNA work, and has been shown to generate a more homogenous representation of targeted positions than the 1240k reagent, and produces data that has been shown to be co-analyzable with shotgun sequencing data (*23*).

We merged paired-end reads into single sequences by requiring an 11-base pair overlap between pairs (allowing for one mismatch if base quality was > 20 and three mismatches if base quality was < 20), which reduces paired sequencing artifacts and ensures that a single molecule is sequenced. Since libraries are sequenced in pools, we used the identifier tags (indices and/or barcodes) to demultiplex raw sequences into individual-specific raw reads in fastq format using a custom tool (available from https://github.com/DReichLab/ADNA-Tools). We stripped adapters and barcodes prior to aligning to the hs37d5 reference sequence which is an hg19 reference sequence with added decoy sequences to reduce alignment artifacts (available from https://www.internationalgenome.org/), using bwa *samse* (v.0.7.15-r1140) with parameters appropriate for ancient DNA where seeding is disabled and gap-opening penalties are modified using: -n 0.01 -o 2 -l 16500. We mapped merged sequences to the RSRS mitochondrial genome (*48*). We removed duplicate molecules from analysis using Picard MarkDuplicates (available from http://broadinstitute.github.io/picard/). We represented the allele at each targeted SNP by randomly sampling overlapping sequences that had a minimum mapping quality ≥ 10 and base quality ≥ 20, excluding two base pairs from the 5’ and 3’ ends to remove damage artifacts.

Ancient DNA authenticity was established using several criteria. We excluded any individuals with a rate of cytosine-to-thymine (C-to-T) substitutions at the terminal nucleotide below 3% (*46*). We computed the ratio of X-to-Y chromosome reads, estimated mismatch rates to the mtDNA consensus sequence (using contamMix v.1.0-12 (*49*)), and estimated X chromosome contamination in males with sufficient coverage (*50, 51*). We also evaluated PCAs for evidence of contamination. Individuals with strong evidence of contaminated data or without at least 20,000 SNPs overlapping targeted positions were excluded. Based on coverage and quality control metrics, we excluded four individuals (I35596, I35606, I37194, and I37195) and retained 19 for genome-wide analysis (**table S1**). Two individuals, I36722 and I36723, were estimated to have ∼1-3% contamination, so we also generated damage-restricted versions of these samples (*52*) and compared the damage-restricted version to the full versions using PCA and *f*_*4*_-statistics. With PCA (**fig. S3**) we observe no shift between versions. All statistics of the form *f*_*4*_(*Full Version, Damage-Restricted Version; European/African, Indigenous American*) were consistent with zero (|Z| < 3), suggesting that any contamination does not systematically bias the allele frequencies toward potential modern contaminant sources. Based on evidence indicating that any contamination present does not affect the major ancestry signal, full versions were used for all population genetics analyses.

### Radiocarbon dates

We report 16 new ^14^C dates on bone material from 15 unique individuals (**table S2**). Two direct dates were generated using accelerator mass spectrometry (AMS) at Beta Analytic and 14 dates were generated in the Pennsylvania State University (PSU) Radiocarbon Laboratory. Beta Analytic carried out conventional gelatin collagen extraction, while at PSU, bone collagen was extracted and purified using a modified Longin method with ultrafiltration (*53*) (>30 kDa gelatin); if collagen yields were low (as was the case for I36726), a modified XAD process (*54*) (XAD amino acids) was used. Carbon and nitrogen isotope ratios were then measured as a quality control measure; all ratios fell between 3.2-3.4, indicating good collagen or amino acid preservation (*53*). All dates reported in the main manuscript were calibrated in OxCal version 4.4 (*55*) using the IntCal20 curve (*56*), and we provide alternate calibrations for the four focal pre-Ceramic Caribbean individuals based on a mixed terrestrial/marine curve and applying a marine reservoir correction in **table S2**. Additional details about ^14^C dating can be found in Supplementary Text.

### Dataset assembly

We merged newly generated genome-wide data from 19 individuals from Samaná who passed quality control screening into a dataset that included genome-wide capture or shotgun sequencing data from 395 ancient Americans (*27, 29, 33-38, 57-59*), 122 present-day individuals (*60-63*), and 4 ancient West Africans (*64, 65*) (**table S15**). To enable the co-analysis of capture- and shotgun-based datasets while reducing technology-specific bias, we restricted all analyses to the Compatibility Panel SNP set which contains 913,708 autosomal SNPs (*24*).

### PCA

We performed Principal Component Analysis (PCA) using the ‘smartpca’ program (version 18720) (*66*). We projected ancient individuals onto the components computed on present-day individuals with ‘lsqproject:YES’ and ‘newshrink: YES’. We ran two PCAs: 1. PCs were computed using 88 present day individuals from three genetically diverse populations: Han, French, and Yoruba. We projected the full and damage-restricted (*_d*) versions of I36722 and I36723 onto these axes, together with three previously published ancient individuals from Argentina (*33*). This analysis was used to assess whether restricting to reads showing characteristic ancient DNA damage altered the placement of these individuals in PCA space. 2. PCs were computed with three genetically distinct ancient American groups with published shotgun sequencing data: *Panama_IsthmoColombian_PreColonial*.*SG (38), BahamasCuba_Ceramic*.*SG (29)*, and *Argentina_BeagleChannel_Yamana_100BP*.*SG (34)*. We projected all ancient individuals and used this PCA to differentiate between pre-Ceramic Age-related and Ceramic Age-related ancestry in the pre-Columbian Caribbean.

### Analysis of shared genomic segments (ROH)

We identified ROH in our ancient dataset using the Python package hapROH (https://test.pypi.org/project/hapROH/) (*39*), which uses 5,008 genomes from the 1000 Genomes project haplotype panel (*67*) as the reference panel. In **table S13**, we group the inferred ROH for the four pre-Ceramic Age individuals from Samaná into four bins: 4-8 cM, 8-12 cM, 12-20 cM, and > 20 cM, and we report the total sum in these bins. To estimate effective community size (*Ne*) we applied a maximum-likelihood inference framework to infer the *Ne* that maximizes the likelihood for ROH lengths observed in a set of individuals. We obtained estimation uncertainties from the likelihood profile: 95% confidence intervals (CI) correspond to values within 1.92 units from the maximum of the log-likelihood function.

### Kinship

We applied a previously described relative detection method (*68*) and identified no close relatives among the individuals from Samaná.

### *f*-statistics

We computed *f*_*4*_-statistics using *qpDstat* (v.1160) from ADMIXTOOLS (*69*) with ‘f4 mode: YES’ and ‘inbreed: YES.’ We computed ‘outgroup’ *f*_*3*_-statistics using *qp3Pop* (v.710) from ADMIXTOOLS (*69*) with ‘inbreed: YES.’ We used pairwise outgroup-*f*3 statistics to visualize shared genetic drift among select ancient American populations. We arranged statistic values into a symmetric matrix and converted to distances as 1/*f*3, such that greater shared drift corresponded to smaller pairwise distances. We then constructed a neighbor-joining tree in R using the ape package and rooted it for visualization with USA_Ancient_Beringian.SG. Negative branch lengths were set to zero before plotting. To improve readability, the tree was ladderized, branch lengths were rescaled for visualization, and terminal branches were equalized so that all tips aligned.

### qpWave

We used *qpWave* v.1610 from ADMIXOOLS (*69*) with ‘allsnps: YES’ and ‘inbreed:NO’ and a core reference (‘right’) set that included Han, Chipewyan, Surui, Piapoco, Karitiana, *BahamasCuba_Ceramic*.*SG, Belize_9600-7300BP, Brazil_LapaDoSanto_older9000BP, Chile_LosRieles_12000BP*.*SG, Peru_Lauricocha_8600BP*, and *Panama_IsthmoColombian_PreColonial*.*SG*. We added populations to this core reference set when carrying out specific tests and include this information in **table S6**.

### qpGraph

We used *qpGraph* (implemented in the admixtools2 package (*70*)) to explore phylogenetic relationships among selected ancient American populations and to evaluate whether simple tree models or models incorporating admixture best fit the data. Analyses were performed using precomputed *f*_2_-statistics in block format generated from the “Compatibility Panel” SNP set. We used an automated graph search strategy implemented in find_graphs() to sample a broad space of candidate admixture graph topologies. For each model class allowing a fixed number of admixture events, we initialized random graphs using random_admixturegraph() and performed multiple independent search replicates. Graph space was explored using a set of topology-altering mutation functions (including subtree pruning and regrafting, leaf swapping, and admixture edge modification), with up to 10,000 generations in the initial search phase and additional refinement steps. For each replicate, multiple starting graphs were optimized in parallel, and up to 10 candidate graphs were retained per run. From each search replicate, we extracted the best-fitting graph (lowest score) and re-estimated parameters using qpgraph() with multiple starting points (numstart = 10). Model fit was evaluated based on the worst residual Z-score across all fitted *f*_*4*_-statistics, with |Z| < 3 considered an acceptable fit. To assess convergence and robustness, we compared graph topologies across replicates by computing graph hashes (using graph_hash()), allowing identification of recurrent solutions.

### Uniparental haplogroups

We determined mtDNA haplogroups for all individuals using .bam files, restricting to reads with MAPQ ≥ 30 and base quality ≥ 20. We constructed a consensus sequence with samtools and bcftools version 1.3.1 using a majority rule and determined the haplogroup with HaploGrep2 (*71*) using Phylotree version 17. We determined Y chromosome haplogroups using sequences mapping to Y chromosome targets, restricting to sequences with MAPQ ≥ 30 and base quality ≥ 30. We called haplogroups by determining the most derived mutation for each individual, using the nomenclature of the International Society of Genetic Genealogy (ISOGG) (http://www.isogg.org) version 14.76 (April 2019).

### Use of Artificial Intelligence (AI)

Generative AI tools (OpenAI’s ChatGPT, GPT-5.4 Thinking) were used during manuscript preparation to assist with text editing and refinement, and to generate or improve R code used in the production of Figs. 1-3 and S1. All outputs generated with AI assistance were critically reviewed, revised, and verified by the authors prior to inclusion.

## Acknowledgements

We thank Kimberly Callan, Trudi Frost, Lora Iliev, Lijun Qiu, and Fatma Zalzala for ancient DNA laboratory work; Matthew Mah, Adam Micco, Nick Patterson, and Gregory Soos for bioinformatics support; Esther Brielle for analytical support; Jeremy Choin and Daniel Tabin for help generating admixture graphs; Iñigo Olalde for help with kinship analysis; Iosif Lazaridis for Y chromosome haplotype determinations; Brendan Culleton for help with calibration of ^14^C dates; and Aisling Kearns for data processing assistance. We appreciate comments from Katherin Nägele. The authors used ChatGPT (OpenAI; GPT-5.4 Thinking) as an AI-assisted tool during preparation of this manuscript. All AI-assisted outputs were reviewed, edited, and verified by the authors. The author-accepted version of this article, that is, the version not reflecting proofreading and editing and formatting changes following the article’s acceptance, is subject to the Howard Hughes Medical Institute (HHMI) Open Access to Publications policy, as HHMI lab heads have previously granted a nonexclusive CC BY 4.0 license to the public and a sublicensable license to HHMI in their research articles. Pursuant to those licenses, the author-accepted manuscript can be made freely available under a CC BY 4.0 license immediately upon publication.

## Funding

National Institutes of Health grant HG012287 (D.R.)

John Templeton Foundation grant 61220 (D.R.)

J.-F. Clin (D.R.)

Allen Discovery Center, a Paul G. Allen Frontiers Group advised program of the Paul G. Allen Family Foundation (D.R.)

Howard Hughes Medical Institute (D.R.)

## Author contributions

Conceptualization: KS AJLB DS DR

Methodology: KS SM NR DR

Investigation: AJLB DS MGA DS

Formal analysis: KS

Visualization: KS

Resources: AJLB DS MGA DS

Supervision: DR AJLB DS

Writing - original draft: KS

Writing - review & editing: KS AJLB DS MGA DS SM NR DR

## Data Availability

The aligned sequences are available through the European Nucleotide Archive under accession number PRJEB73283. Genotype data used in analysis are available at https://reich.hms.harvard.edu/datasets. Any other relevant data are available from the corresponding authors upon reasonable request.

## Code Availability

The code used for data processing in this study are available from https://github.com/DReichLab/ADNA-Tools. The ADMIXTOOLS code for used for population genetic analyses are available from https://github.com/DReichLab/AdmixTools.

