## Supplementary Materials for "First Lithic Age Caribbean genomes document pre-Ceramic genetic continuity and affinities to Central America and northern South America"

Kendra Sirak *et al.*

### The PDF file includes:

Supplementary Text
Figs. S1 to S3
References

### Other Supplementary Material for this manuscript includes the following:

Tables S1 to S15

**Supplementary Text**

### Archaeological Overview

The Guahayona Institute, presided by co-author Daniel Shelley, and its team of archaeologists, directed by co-author Adolfo López, began excavating in the Samaná Peninsula of Hispaniola in 2018. Excavations started with the Ceramic Age site of El Francés. However, based on Shelley's hypothesis of links between Lithic Age Casimiroids and the Ceramic Age 'Ciguayo' inhabitants of El Francés (*1*) who lived on and around the Samaná Peninsula and who differed in language and customs from the people referred to as 'Taíno' who occupied other parts of Hispaniola (*2, 3*), the team prospected for Lithic Age and later sites nearby and discovered a complex of cave and rock shelter sites about 8 km north of El Francés in the Monumento Natural Cabo Samaná. The skeletal samples we analyze in this work, excavated in 2022-2023, come from these sites as well as the neighboring site of Playa Madama, located ~5 km north.

### Skeletal material

The individuals analyzed in this study were recovered in a range of burial positions, reflecting diverse mortuary practices. Notably, the four individuals with pre-Ceramic Age-associated genetic signatures (I36726, I36721, I36722, and I36723) were all interred in a consistent burial position that is qualitatively distinct from that observed in individuals with Ceramic Age-associated genetic signatures.

Individual I36726 (CF2), whose bone material yielded a pre-Ceramic Age-associated genetic signature and whose direct $^{14}$C dates fall within the Lithic Age 4411-4158 calBP (3870±30 BP, PSUAMS-14458) was found at Cueva Funeraria de Daniel; the archaeologists obtained radiocarbon dates from associated charcoal samples from the same burial pit that are consistent with the direct radiocarbon date after adjusting for the possibility of a one to two century marine reservoir effect for the bone-based date (**table S2**). This individual was found in a supine position, with arms extended down the sides and pressed against the ribs, the legs straight and close together, and the skull wedged down between the shoulder blades, which suggests that the body was wrapped at the time of burial, perhaps in some kind of shroud.

The archaeologists found the same positioning in the excavation of three individuals I36721 (AD2), I36722 (AD3), and I36723 (AD7) from the Abrigo Daniel, whose bone material also yielded pre-Ceramic Age-associated genetic signatures. The identical burial style of these individuals indicates that the bodies were intentionally positioned this way. Like I36726, I36721 and I36722 were also subject to some uniform pressure that fixed them firmly in position at the time of burial, likely some kind of funerary wrapping. The archaeologists found one other individual buried in this position at Abrigo Daniel, from whose bones no genetic material was obtained (I37188/AD5). Direct dates on bone places I36721 as having lived 4079-3845 calBP (3630 ± 30 BP, Beta-765015) and I36723 as having lived 4150-3933 calBP (3710 ± 30 BP, Beta-765018). Charcoal samples collected near these individuals (and I37622) at Abrigo Daniel yielded the following dates: 5285-4886 calBP (4450 ± 30 BP, Beta-638467) and 5313-5051 calBP (4530 ± 30 BP, Beta-638468).

The burial style of the individuals with pre-Ceramic Age-associated genetic signatures at the Samaná Peninsula sites, three of whom have a directly calibrated radiocarbon date which falls in the Lithic Age, is different in kind from that of the individuals with Ceramic Age-associated genetic signatures and directly calibrated radiocarbon dates which fall in the Ceramic Age.

### Radiocarbon dating

We generated new direct radiocarbon ($^{14}C$) dates for 15 unique individuals using bone material. In **table S2**, we report the conventional radiocarbon age, isotopic information, and provide different calibrations of the conventional radiocarbon age.

Radiocarbon dates from human bone from island contexts are often corrected for a marine radiocarbon reservoir effect (dR), and, when possible, calibrated using isotopic information from ancient individuals to provide guidance about the proportion of an individual's diet composed of marine resources. Assessing the proportion of an individual's diet composed of marine resources in the Caribbean is challenging, because fish and mollusks from coral reef environments have a $C_4$-like dietary signature (average $d^{13}C$ of about -10 per mil), while terrestrial foods are mostly $C_3$ (average $d^{13}C$ of about -23 per mil) (*4*). However, maize (which probably arrived in the Caribbean in pre-Ceramic times (*5*)) has a $C_4$ signature of about -10 per mil (i.e., it looks marine), while *Codakia orbicularis* clams have a -23 per mil signature due to chemoautotrophic sulfur bacteria synthesizing food in their gills (i.e., they look terrestrial). Even in the pre-Ceramic Age before agriculture became widespread, the degree of correction needed based on diet is complex. As the success of fishing versus terrestrial foraging is variable and there is evidence of fairly rapid local resource depletion in the Caribbean even at low population densities (see refs. (*6, 7*)), it is even more difficult to accurately evaluate the contribution of marine resources to an individual's diet. Consistent with previous publications reporting ancient DNA from pre-contact inhabitants of the Caribbean (*8, 9*) we report dates in the main manuscript using the IntCal20 curve without a marine reservoir correction so as to not introduce additional systematic error based on dietary variation.

However, we have evidence that at least some of our direct dates on bone are producing calibrated calendar dates that may be older by on the order of a century or two than the true calendar dates, suggesting that a marine reservoir effect correction may be needed. In particular, for individual I36726 (CF2), we have a date both from the same bone from which we extracted DNA (3870 ± 30 BP, PSUAMS-14458), and from charcoal from the same burial pit (3720 ± 30 BP, Beta-638474), and the bone date is 150 uncalibrated years older ($P=0.0004$ for being significantly older). Therefore, in **table S2**, we also provide dates calibrated using a mixed marine (Marine20) and terrestrial (IntCal20) calibration curve based on different proportions of marine contribution to the diet, as well as a marine reservoir correction. We base our marine reservoir correction (dR) on data from ref. (*10*) who provide dR values for the Dominican Republic and in particular, for Samaná Bay. We compute dR as the weighted average of the two *Argopecten* estimates (excluding the outlier *Codakia* from Cap-Haitien) and obtain a dR of -319 ± 21 $^{14}C$ yr. To calibrate conventional radiocarbon dates accounting for a 25 ± 10% or a 50 ± 10% marine contribution to diet, we use a mix of the IntCal20 and Marine20 curves, coding for either a 25 ± 10% or 50 ± 10% proportion of the Marine20 curve.

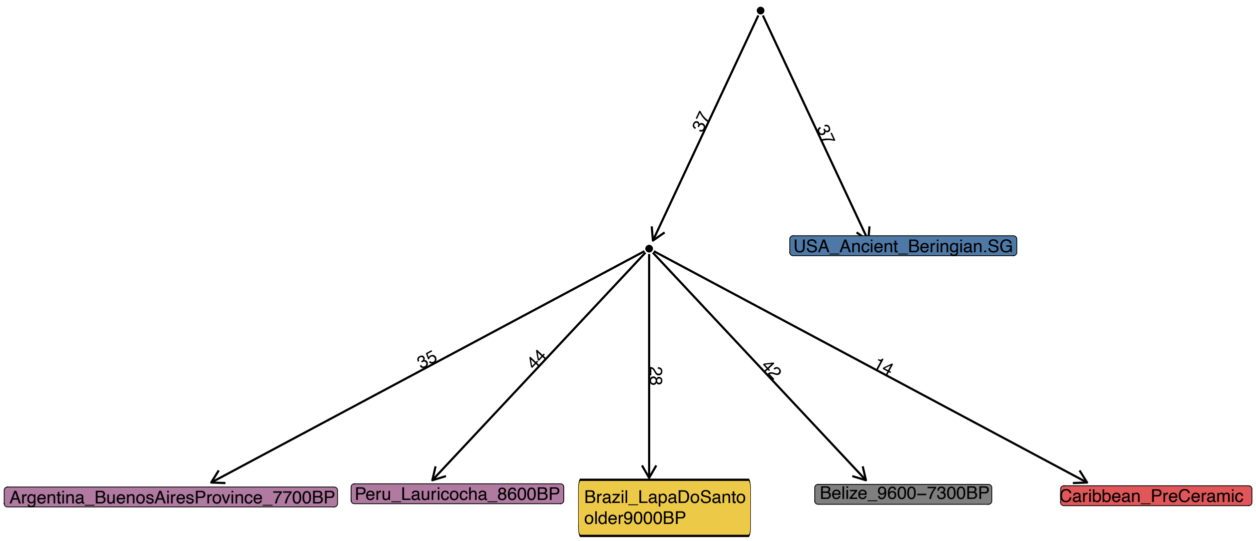


**Fig. S1. Simple admixture graph with no admixture events for six populations representing deep structure in the Americas.** We show a *qpGraph* model including six populations – *USA_Ancient_Beringian.SG, Argentina_BuenosAiresProvince_7700BP, Brazil_LapaDoSanto_older9000BP, Peru_Lauricocha_8600BP, Belize_9600–7300BP, and Caribbean_PreCeramic* – that provides an adequate fit without requiring admixture (score = 6.9; $|Z|_{max}$ = 2.4). Very short branches (drift < 0.001) are collapsed for visualization. In this reduced model, *Caribbean_PreCeramic* is placed as a distinct lineage within SNA-related variation. This graph captures broad-scale relationships among deeply divergent populations but does not include populations from Central America or northern South America that may share finer-scale ancestry with *Caribbean_PreCeramic*. As a result, the topology should be interpreted as a simplified approximation of population history rather than a uniquely supported model.

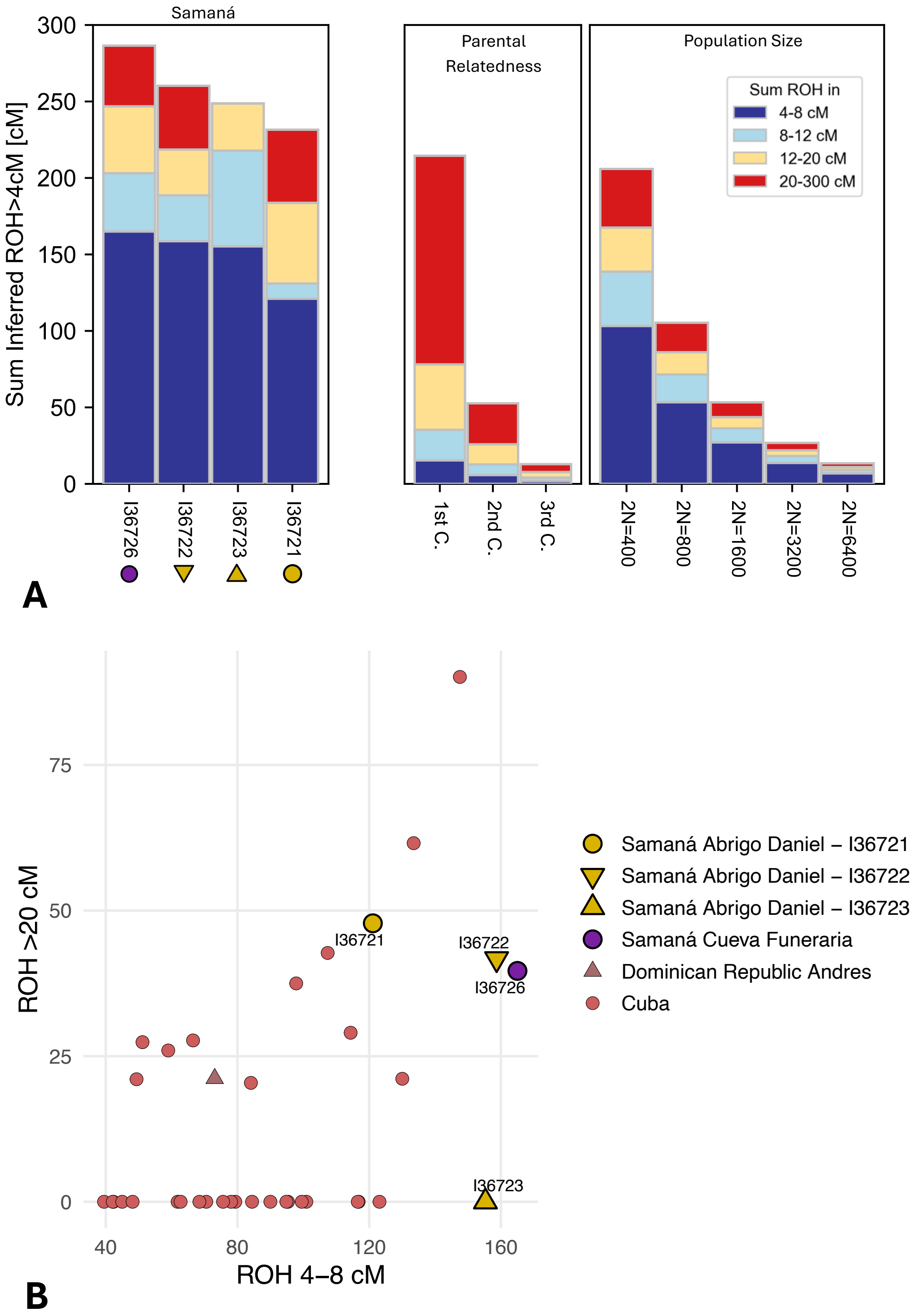


**Fig. S2. Analysis of ROH in Samaná and comparison with ROH elsewhere in the pre-Ceramic Age Caribbean.** (**A**) Distribution of ROH for four individuals from Samaná. On the left, we depict the total sums of ROH that fall into four length bins: 4-8cM (dark blue), 8-12cM (light blue), 12-20cM (yellow), and >20cM (red) for three individuals with pre-Ceramic Age genetic signatures from Abrigo Daniel and one from Cueva Funeraria. We show analytical expectations based on parental relatedness (C. stands for cousin) and population size on the right. (**B**) Long (>20cM) ROH plotted against short ROH (4-8cm) for all previously published and new pre-Ceramic Age Caribbean individuals with sufficient coverage.

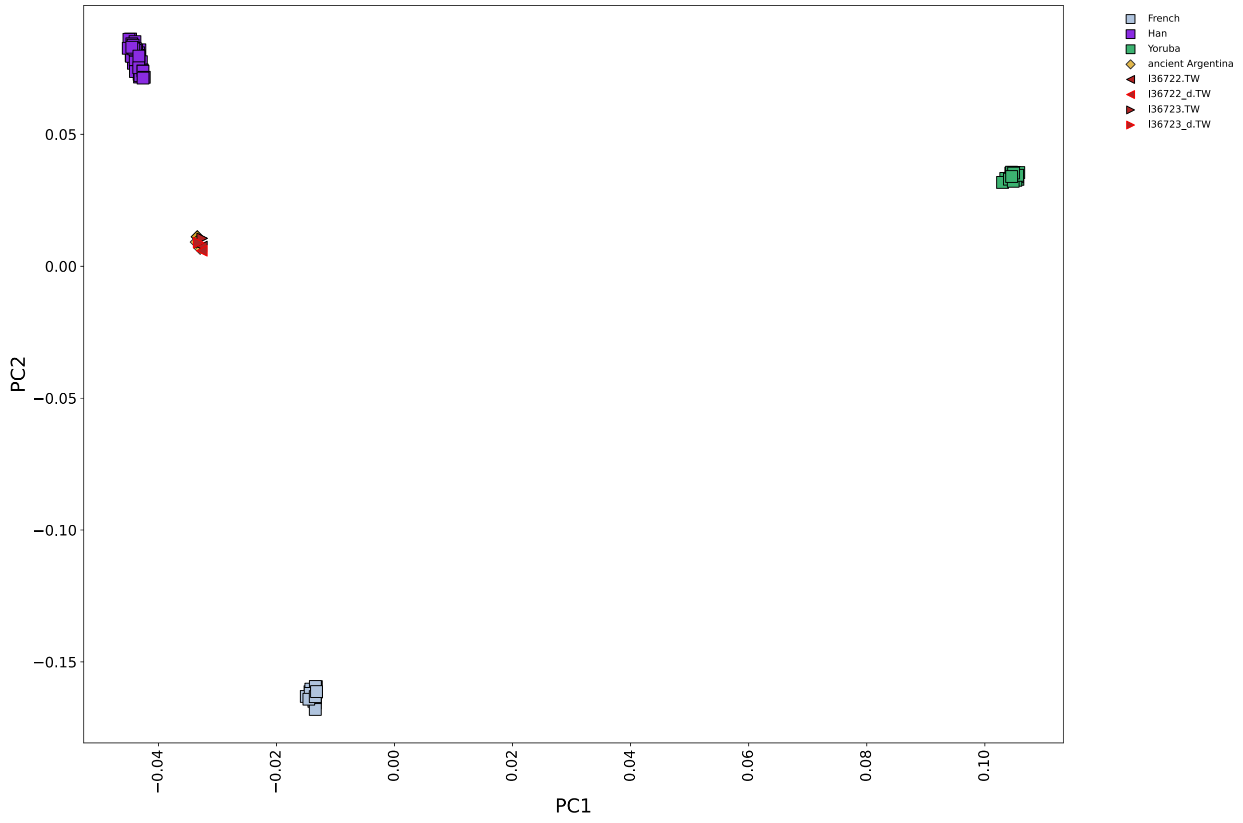


**Fig. S3. Worldwide PCA showing damage-restricted (_*d*) and full versions of I36722 and I36723 projected onto principal components computed from present-day French, Han, and Yoruba individuals.** Ancient Argentinean individuals (*11*) are shown for reference. The damage-restricted (_*d*) and full versions of I36722 and I36723 occupy the same positions in PCA space, indicating that restricting to damaged reads does not alter their broad genetic placement. This supports the conclusion that the major ancestry signal in these individuals is not impacted by any contamination.
